# Construction of the ETECFinder database for the characterisation of enterotoxigenic *Escherichia coli* (ETEC) and revision of the *VirulenceFinder* webtool at the CGE website

**DOI:** 10.1101/2023.04.11.536449

**Authors:** Flemming Scheutz, Camilla Hald Nielsen, Astrid von Mentzer

**Affiliations:** The International *Escherichia* and *Klebsiella* Centre, Statens Serum Institut, Denmark; Department of Bacteria, Parasites and Fungi, Statens Serum Institut, Denmark; Department of Microbiology and Immunology, Institute of Biomedicine, Sahlgrenska Academy, University of Gothenburg, Sweden; Parasites and Microbes Programme, Wellcome Sanger Institute, Hinxton, United Kingdom

## Abstract

The identification of pathogens is essential for effective surveillance and outbreak detection, which has been facilitated by the decreasing cost of whole-genome sequencing (WGS). However, extracting relevant virulence genes from WGS data remains a challenge. In this study, we developed a web-based tool to predict virulence-associated genes in enterotoxigenic *Escherichia coli* (ETEC), including heat-labile toxin (LT) genes (*eltA* and *eltB*), heat-stable enterotoxin (ST) genes (*est*) for both human and animal, enterotoxin (*espC*), colonization factors CS1 through 30, F4, F5, F6, F17, F18, and F41, and toxigenic invasion and adherence loci (*tia, tibAC, etpBAC, eatA, yghJ*, and *tleA*).

To construct the database, we revised the existing ETEC nomenclature and used the VirulenceFinder webtool at the CGE website (https://cge.cbs.dtu.dk/services/VirulenceFinder/). The database was tested on 1,083 preassembled ETEC genomes and the ETEC reference sequence for strain H10407. We added 455 new alleles, replaced or renamed 50 alleles, and removed two.

Overall, our tool has the potential to greatly facilitate ETEC identification and improve the accuracy of WGS analysis. The revised nomenclature and expanded gene repertoire provide a better understanding of the genetic diversity of ETEC. Additionally, the user-friendly interface makes it accessible to users with limited bioinformatics experience.

## INTRODUCTION

Enterotoxigenic *Escherichia coli* (ETEC) causes hundreds of millions of cases of infectious diarrhea each year, mainly in developing countries (1) and is ranked number 7 on the global burden of 31 food-borne hazards (2). Globally, one in ten child deaths in children under the age of five is a result of diarrheal disease, resulting in 800,000 deaths annually (3). ETEC is also the most common cause of *Escherichia coli* (*E. coli*) diarrhea in farm animals (mainly cattle, pigs, and sheep) (4). The European Food Safety Authority (EFSA) reports that multidrug-resistant (MDR) *E. coli* is considered an important hazard to public health. The indication associated with most antimicrobial use is post-weaning diarrhea (PWD) caused by ETEC (5) and recent studies show that there is an increase in antibiotic resistance seen in ETEC infecting humans (6–9). The presence of multidrug-resistant pathogenic bacteria, including ETEC, in raw vegetables and minimally-processed fresh vegetables, is a significant public health concern (10).

ETEC bacteria colonize the small intestine via colonization factors (CF), also known as coli surface antigens (CS) or fimbriae (F), and the secretion of enterotoxins, heat-labile toxin (LT) and/or heat-stable toxin (ST), alters the epithelial cell systems (11, 12) resulting in secretory diarrhea and dehydration. Infections can become lethal as a result of severe dehydration and electrolyte imbalance (4) and is a huge problem in many pig herds as well as in geographical regions where access to clean water is limited.

At least 28 antigenically distinct CFs and 30 LT types have been identified and characterized in human ETEC (13, 14) and GitHub - avonm/ETEC_toxin_variants_db and six CFs in ETEC isolated from animals. The enterotoxins LT and ST can be further sub-grouped: LTI comprises LTh and LTp (nomenclature is based on human or porcine origin). At least 28 LT types have been identified (13). ST may be classified into two major genotypes, *i.e.*, STa and STb (15). Two variants of STa are very similar in structure and function and grouped into STh (16) and STp (15) based on which host they were originally found in. STh consists of 18 amino acids, and STp of 19 amino acids (17). However, both ST types are associated with infections in humans (15) and animals. STa is also found in *Vibrio cholerae* non-O1 and O1, *Vibrio mimicus, Yersinia enterocolitica, Klebsiella pneumonia* and *Citrobacter freundii* (17, 18).

The cost of whole-genome sequencing (WGS) has decreased over the last decades, making the technology accessible to routine clinical and microbiological laboratories. Even though WGS provides detailed information that could enable diagnostics and typing based on the information obtained from the bacterial genome, the challenge is to extract the relevant information from the large amount of sequence data that is generated by this technique. Thus, it is important that the data can be interpreted by physicians, veterinarians and public health professionals, and to achieve this, an ETECFinder database has been constructed and incorporated into the pre-existing webtool, *VirulenceFinder*, provided by the Center for Genomic Epidemiology (CGE) (www.genomicepidemiology.org). The webtool is user-friendly and allows for rapid analysis of the obtained WGS data and extraction of relevant information, useful for diagnostics, surveillance, and outbreak investigations. The ETECFinder contains every known ST, LT-gene, CFs related to human and animal disease and other genes known to be involved in ETEC virulence/pathogenicity.

There is no perfect test for ETEC: detection of colonization factors is impractical because of their great number and heterogeneity. But ETEC infections pose a major threat to global health in a One Health environment. Rapid diagnostics and accurate classification are of great importance in order to limit the spread of and prevent outbreaks. Diagnostics of ETEC isolates are seldom done in a laboratory, but based on the patient’s or animal’s history and symptoms, thus making monitoring of ETEC very difficult. By expanding the existing *VirulenceFinder* tool and database, we hope to contribute to a more refined and accurate detection of virulence factors related to ETEC infections in both humans and animals.

## MATERIALS AND METHODS

### Construction of the *VirulenceFinder* database

The relevant genes and their sequences were compared using BLASTn against the NCBI nucleotide database (https://www.ncbi.nlm.nih.gov/nuccore/) to identify potential matches. Sequences with open reading frames (ORFs) were curated and validated for the presence of only the four nucleotides ATCG. The validated sequences were compared using BioNumerics software (v8.1), and a reference sequence was selected for each allele. The first validated sequence to enter the database was designated as the reference sequence.

To collect alleles, sequences from various papers were compared using BLASTn and alleles that matched 80% sequence identity and total length were included in the database. Partial genes were excluded from the database to ensure completeness of sequences. The database is stored as a text file in FASTA format with a unique identifier for each reference sequence followed by its sequence.

### Update of the *VirulenceFinder* database

The original *VirulenceFinder* database already contained 14 genes (original designations: *cfa_c, cofA, f17-A, f17-G, fanA, fasA, fedA, fedF, K88ab, fimF41, lngA, ltcA, sta1 and stb*) (see **Supplementary Table 1**) associated with ETEC. To expand the database, the full *loci* for CFs CFA/I, CS8 and CS21 as well as animal-specific fimbriae F4, F5, F6, F17, F18, and Fim41, were added. The genes and *loci* were revised, updated, and added (see **Supplementary Table 1**). The original designation *ltcA*, where “c” stands for chicken (19) was deleted.

The LT toxins presented a challenge when using nucleotide sequences. The original definitions of LT types were based on the specific combination of the A and B subunits (**Supplementary Table 2**). For instance, LTIh-3 and LTIh-5 have identical A subunits, as do LTIp-1, LTIh-4 and LTIh-6. The same applies for the B subunit of LTIh-9, LTIh-11 and LTIh19; LTIh-17, LTIh-20, LTIh-29 and LTIh-30; LTIh-3 and LTIh-8; LTIh-2, LTIh-7, LTIh-15, LTIh-16 and LTIh-22; LTIh-1a, LTIh-1b, LTIh-10, LTIh-12, LTIh-13, LTIh-18, LTIh-21, LTIh-23, LTIh-24, LTIh-25, LTIh-26, LTIh-27 and LTIh-28 respectively. However, the reference sequences submitted to NCBI vary in whether or not the nucleotide sequences encoding the holotoxin include the overlap of the stop codon for the sequence encoding the A subunit and the start codon encoding the B subunit. Sequences *eltI*-2, *eltI*-17, *eltI*-18, *eltI*-19, *eltI*-20, *eltI*-21, *eltI*-22, *eltI*-23, *eltI*-24, *eltI*-25, *eltI*-26, *eltI*-27 and *eltI*-28 have no overlap and are 1,152 bp whereas the remaining *eltI* sequences (*eltI*-1a, *eltI*-1b, *eltI*-3, *eltI*-5, *eltI*-6, *eltI*-7, *eltI*-8, *eltI*-9, *eltI*-10, *eltI*-11, *eltI*-12,1 *eltI*-3, *eltI*-14, *eltI*-15, *eltI*-16, *eltI*-29 and *eltI*-30) are 1,148 bp. Due to the high sequence identity between the *eltI* (*elth*) sequences (>98.5%), the four bp difference may result in a wrong *eltI* type for tested sequences using the CGE *VirulenceFinder* tool. As an illustration, the *eltI*-1 (1,148 bp with an overlap) has a non-synonymous SNP at position 88 compared to *eltI*-25 (1,152 bp with no overlap). To address this difference, we removed the “TGAA” nucleotides from the *eltI* reference sequences (*eltI*-2, *eltI*-17, *eltI*-18, *eltI*-19, *eltI*-20, *eltI*-21, *eltI*-22, *eltI*-23, *eltI*-24, *eltI*-25, *eltI*-26, *eltI*-27 and *eltI*-28) in our database. The final revision now includes the *eltI*AB sequence that encodes the holotoxin for 30 human LTIh types (LTIh-1 through LTIh-30), with two LTIh-1 alleles, and one porcine LTIp type (LTIp-1). Additionally, we identified and analyzed 15 holotoxin sequences for LTII (*eltII*AB) and added them to the database. These sequences consist of one LTII-a, two LTII-b, nine LTII-c, two LTII-d and one LTII-e.

To account for the variations in the porcine variant of STa1, we replaced the two *sta1* alleles with 15 *estap* alleles. For the human variant of STa2 and STa3 we added four *estah* alleles. Furthermore, we renamed three *stb* alleles for the porcine variant STb1 as *estb*, and added one variant of ST2b to the database (see **Supplementary Table 1**).

### Creation of the ETECFinder database

We used a beta version of the *VirulenceFinder* database, which includes the ETECFinder database (ETEC-related genes), to search for virulence genes in 1,083 preassembled *E. coli* genomes. The ETECFinder database is designed to perform a genotypic detection of ETEC virulence genes, and all genes and alleles in the database are identified with a GenBank accession number or identifier (ID). To simplify the LT typing, we concatenated the *eltA* and *eltB* sequences and removed the overlapping four bases present in a subset of the LT types. We then analyzed the nucleotide sequences that encodes the LT holotoxin. Additionally, we ran the 1,083 genomes through the CGE webtool *SerotypeFinder*, CGE Server (dtu.dk).

Genomes were defined as ETEC if they were positive for any combination of the genes *estah*, *estap*, *estb*, *eltI*AB or *eltII*AB. Additionally, genomes were classified as ExPEC_JJ_ if they were positive for two or more of *papAH* and/or *papC* (P fimbriae), *sfa/focDE* (S and F1C fimbriae), *afa/draBC* (Dr-binding adhesins), *iutA* (aerobactin siderophore system), and *kpsM* II (group 2 capsules) (20), and as UPEC_HM_ if they were positive for two or more of *chuA* (heme uptake), *fyuA* (yersiniabactin siderophore system), *vat* (vacuolating toxin), and *yfcV* (adhesin) (21).

### Validation of the database

A collection of 1,083 ETEC isolates were selected for sequencing as part of other studies with the aim of representing the global diversity of ETEC from 57 different countries in Asia, Africa, and North, Central and South America between 1980 and 2011. Isolates were chosen based on their host and virulence profile to cover the range of ETEC diversity. This includes isolates collected from adults (both indigenous and travelers), children under 5 years of age, and farm animals infected with ETEC. A subset of the clinical samples was collected from asymptomatic individuals. The isolates were screened for toxins using PCR and a subset of CFs using dot-blot with available antibodies. A subset of the genomes was manually analyzed for virulence profile, while DNA extraction, sequencing, and assembly of the additional 721 genomes followed the same protocol as described in von Mentzer *et al.* (22). An additional level of quality control of the reads and assemblies was performed to ensure high-quality sequencing data. FASTQC v0.11.8 (<u>Babraham Bioinformatics - FastQC A Quality Control tool for High Throughput Sequence Data</u>) and MultiQC (23) were used to investigate read quality and GC content (between 49% and 51%). Kraken/bracken was used to identify potential contamination in combination with assembly statistics such as species abundance (>65% *E. coli*), the total number of bases (4.5-6 MB), the total number of contigs (<300), and the N50 value was used to further assess the quality of the assemblies. Finally, the annotated sequences for ETEC reference strain H10407 (chromosome, Acc. No. FN649414, and plasmids FN649415.1_p52, FN649415.1_p58, FN649415.1_p666 and FN649415.1_p948) were tested using the revised database.

## RESULTS

Including original, renamed, replaced and new genes, the beta ETECFinder database contains 524 alleles representing 38 *loci;* the gene name, the number of new alleles, and the CFs or protein names are listed in **Supplementary Table 3**. These alleles were added to and/or changed in the existing database at the CGE website.

### Updated nomenclature for enterotoxins

When constructing the ETECFinder database, a search for the various ST- and LT types was conducted, which revealed identical genes but with different nomenclature. To ensure that identical gene sequences do not have multiple names, a revised, new nomenclature is proposed and listed in **Table 1**, along with the present nomenclature and accession numbers of the reference genes.

**Table 1.**
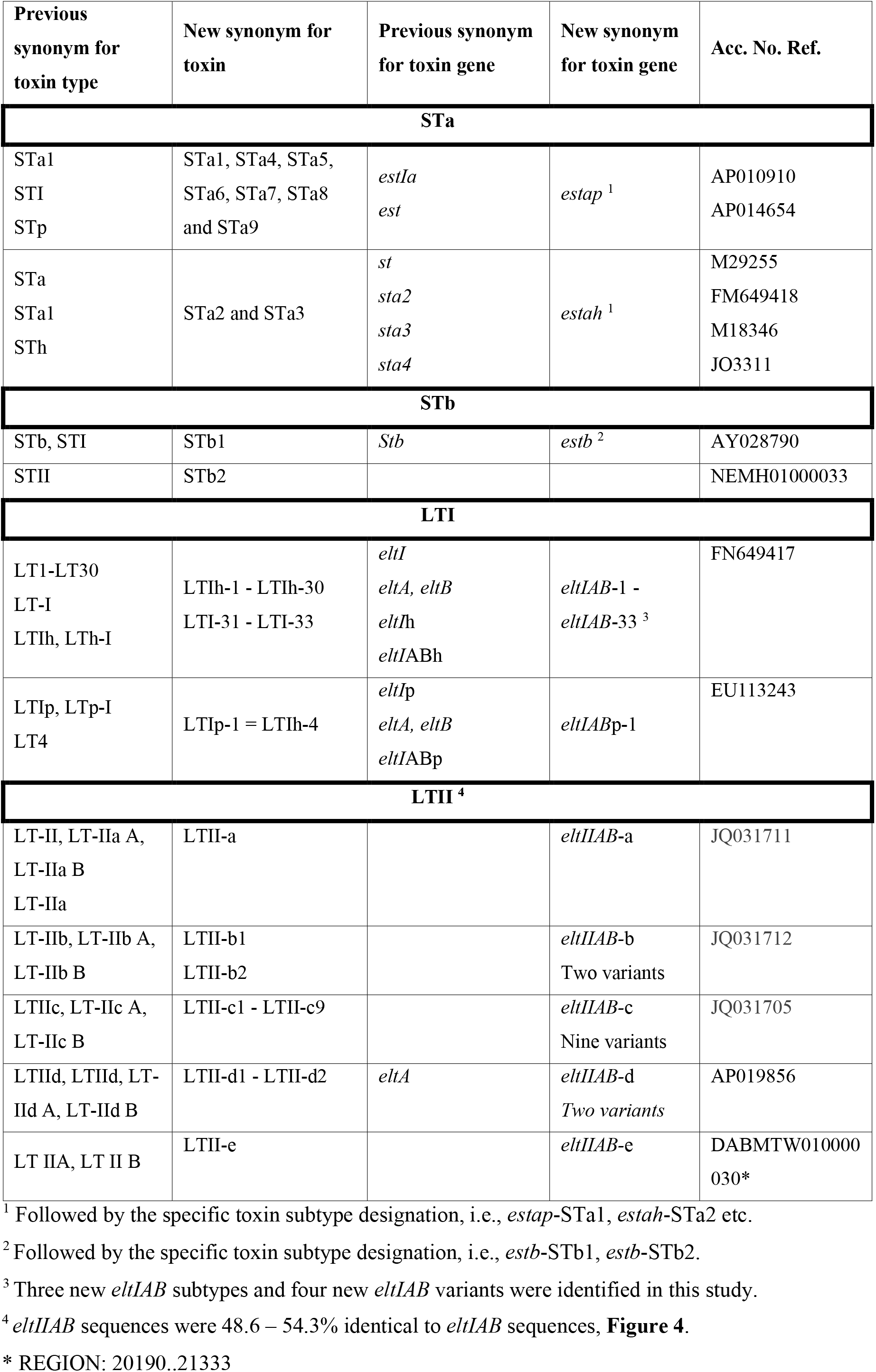
Nomenclature of LT and ST toxins and genes.

As a result of the new nomenclature, the existing *VirulenceFinder* webtool had to be revised. The revision led to changes in the nomenclature of *est* and *elt* genes and the number of alleles. Some ETEC virulence genes were previously found in the *VirulenceFinder* database, and these represented different CF, ST and LT types. However, the database did not contain genes from the entire fimbriae but merely one or two genes. As a result of this study, genes from the entire CF *loci* have been added to the database. Changes in the *VirulenceFinder* database are listed in **Supplementary Table 1,** and new alleles in **Supplementary Table 3**.

### Implementation of ETECFinder

The ETECFinder database comprising all known ETEC virulence genes is now a part of the VirulenceFinder tool, which, together with the notes file, can be downloaded from the CGE website (<u>genomicepidemiology / virulencefinder_db — Bitbucket</u>). Currently, VirulenceFinder can either take reads or assembled (complete or partly assembled) as input. Because the CGE webtools only allows for one sequence at a time, a script for batch analyzes was kindly provided by bioinformatician Maja Weiss (CGE) so that batch analyzes could be performed on the SSI server. Firstly, raw data generated from sequenced and assembled bacterial genomes were used as input. By performing a BLAST search of the genome against the database, the closest matching alleles are identified, and a CF and toxin profile is determined. The CF profile is based on the various alleles found. The short output format includes the identified best-matching ETEC allele in the database. An additional extended output includes the nucleotide sequence of the ETEC alleles identified. For a few selected outputs, both types of input data - reads and assembled genomes - were analyzed and compared. **Figure 1** shows the output using the preassembled genome of sequence 31919_4_289, and Figure 2 shows the output using FASTQ files of the same strain in the revised database. Using the revised database on this sequence, an additional 12 alleles representing six genes were identified, and *ltc* and sta1 were identified as *eltIAB*-23 and *estap*-STa1, respectively (data not shown). In this particular run, the genes *fdeC and hha* were not identified using the FASTQ files. However, *csgA, csmC, csmD, iss* (two variants), and *traT* (three variants) were identified using FASTQ files, **Figure 1 and Figure 2**.

**Figure 1.**
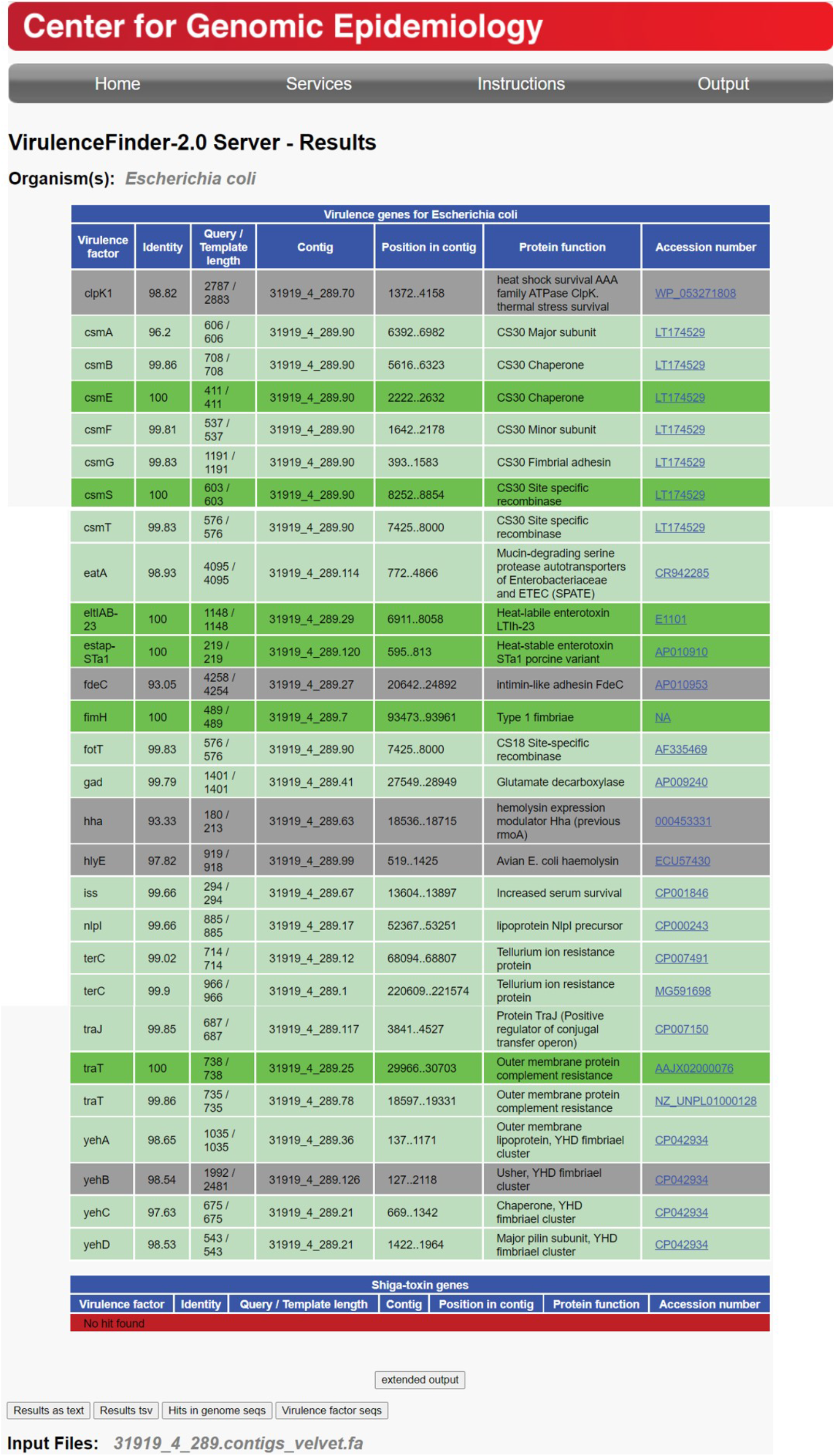
ETECFinder results for an Enterotoxigenic *Escherichia coli* isolate in the short output format using the revised VirulenceFinder database with a preassembled genome of an Enterotoxigenic *Escherichia coli.* Shown are the names of the best-matching allele in the VirulenceFinder, the percentage of nucleotides that are identical to the best-matching allele in the database and the corresponding sequence in the genome (% identity), the length of the alignment between the best-matching allele in the database and the corresponding sequence in the genome (also called the high-scoring segment pair [HSP]), the length of the best-matching allele in the database, the name and function of the best-matching allele, and an LT type. Color indications: The dark green color indicates a perfect match for a given gene. The %Identity is 100, and the sequence in the genome covers the entire length of the virulence gene in the database. The light green color indicates a warning due to a non-perfect match, %ID < 100%, HSP length = virulence gene length. The grey color indicates a warning due to a non-perfect match, HSP length is shorter than the virulence gene length, %ID = 100%. The red color indicates that no virulence gene with a match over the given threshold was found. See https://cge.cbs.dtu.dk/services/VirulenceFinder/output.php.

**Figure 2.**
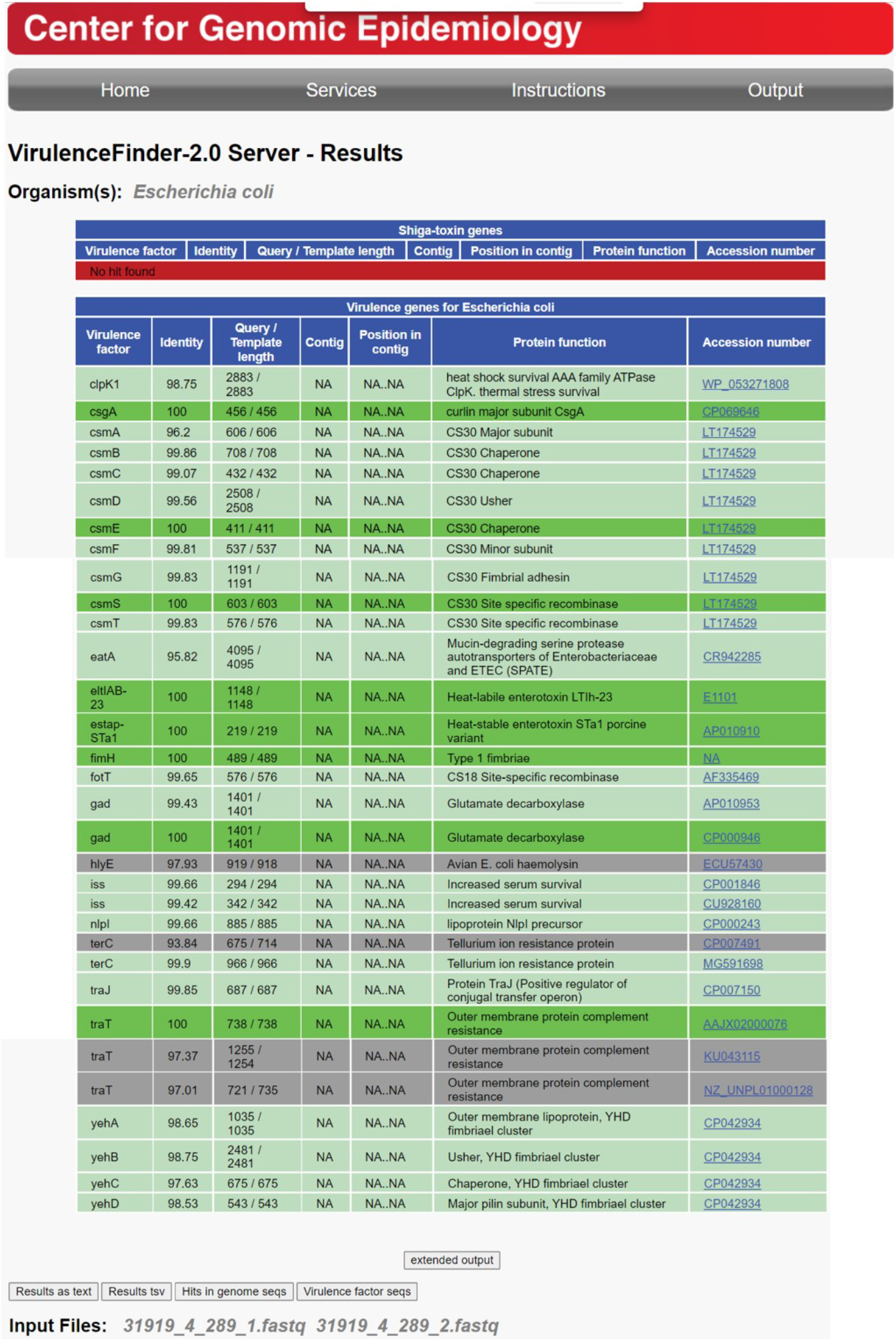
ETECFinder results for an enterotoxigenic *Escherichia coli* isolate in the short output format using the revised VirulenceFinder database with FASTQ files from the same enterotoxigenic *Escherichia coli* as in Figure 1.

Multiple new virulence gene alleles identified

### Verification of the ETECFinder

The revised database was tested on a collection of 1,083 preassembled genomes (sequenced by Astrid von Mentzer) to predict the virulence profile and compared to previous bioinformatics analyzes. The threshold for a positive hit was set at 90% sequence identity and minimum length at 60%. This resulted in 890 ETEC matches (*i.e*., enterotoxin-positive *E. coli*) and 193 non-ETEC matches (*i.e*., enterotoxin-negative *E. coli*). One ETEC sequence also qualified as UPEC_HM_ (ETEC-UPEC_HM_) and four as ExPEC_JJ_ (ETEC-ExPEC_JJ_). Two sequences qualified as UPEC_HM_, 15 as ExPEC_JJ_, and 13 as ExPEC_JJ_-UPEC

### Enterotoxins

In total, 890 (82%) genomes were toxin positive, and in 193 (18%) genomes, no toxin genes were detected. The most prevalent toxin profiles were STh-only (193/1083; 18%), LT-only (184/1083; 17%), LT+STh (157/1083; 14%), STp-only (100/1083; 9%). Additional toxin profiles with LT in combination with other toxins were LT+ STh (157/1083; 14%), LT+STp (109/1083; 10%), LT+STb (107/1083; 10%), LT+STp+STb (18/1083; 1.7%) and LT+STh+STp (8/1083; 0.7%). A new variant of STb was discovered in one genome. Thirteen genomes had a combination of STa1 and STb1. STa1 and STa3 were only found in combination with LT18 in eight genomes of serotype O148:H28. The most common combination of LT and ST type was LTIp and STb1 (107/1083; 10%), followed by LT30 and STa3 (90/1083; 8%) (**Supplementary Table 4)**. Three new LT types were discovered where LT32 was identified in 94 genomes (51 alone and 15 together with STa1), followed by LT33 (12 alone and 15 with STa1) and LT31 with STa1 (1). Additional new variants designated LT15b (11/1083), LT12b (2/1083), LT18b with STa1 (6/1083) and LT18c with STa1 (1/1083) were found in the analyzed genomes (**Figure 3)**.

**Figure 3.**
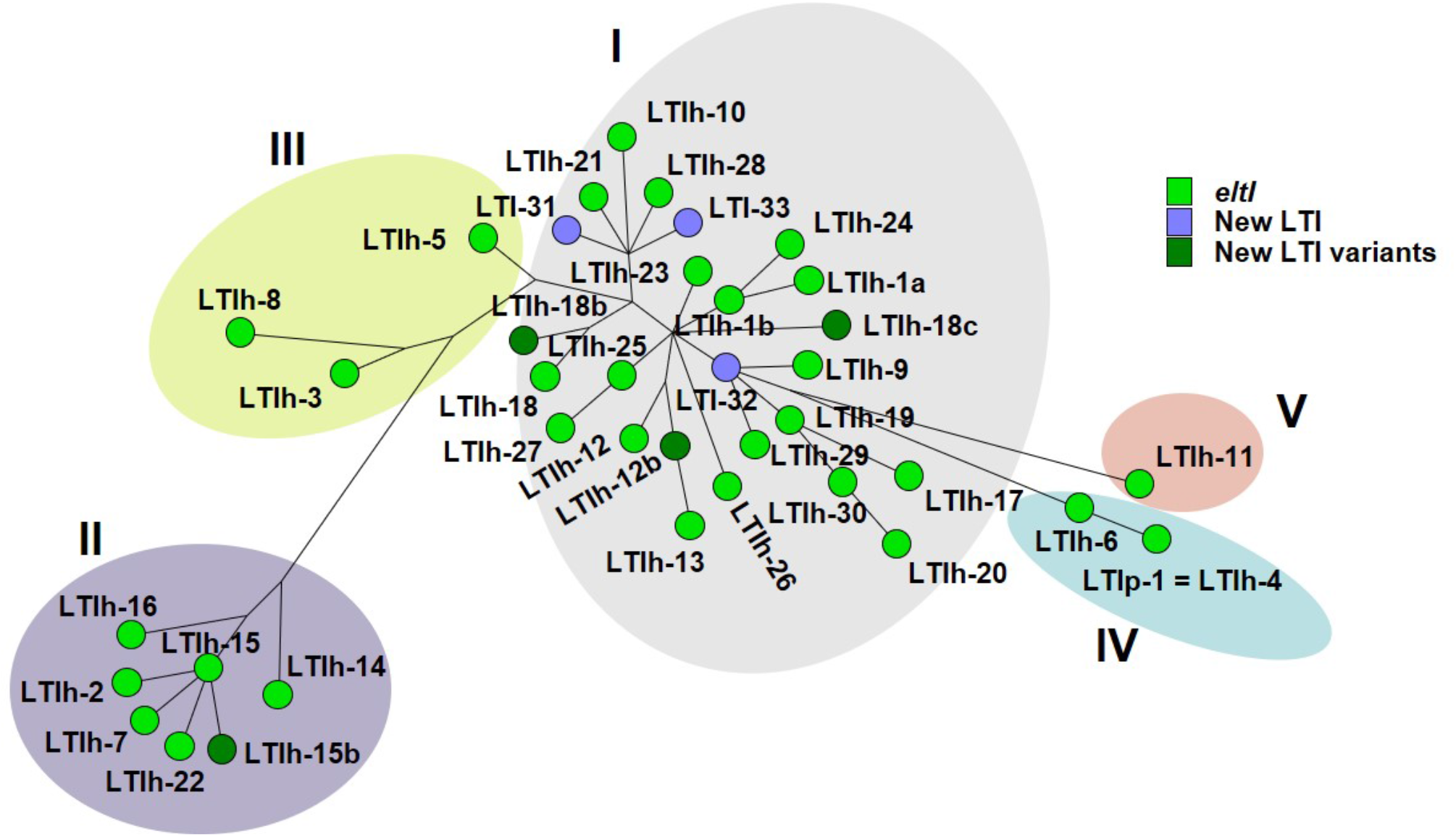
Maximum parsimony tree of 31 LTI reference nucleotide sequences, three new LTI types (LTI-31, LTI-32 and LTI-33) and four new LT variants) LTIh-12b, LTIh-15b, LTIh-18b and LTIh-18c) identified in 890 ETEC genomes analyzed with the revised VirulenceFinder database. The colors and groups I – V correspond to the designation and colors in Joffré *et al.* (13) where concatenated protein sequences were presented. Four nucleotides “TGAA” were removed from the NCBI-submitted sequences as described in the text.

**Figure 4.**
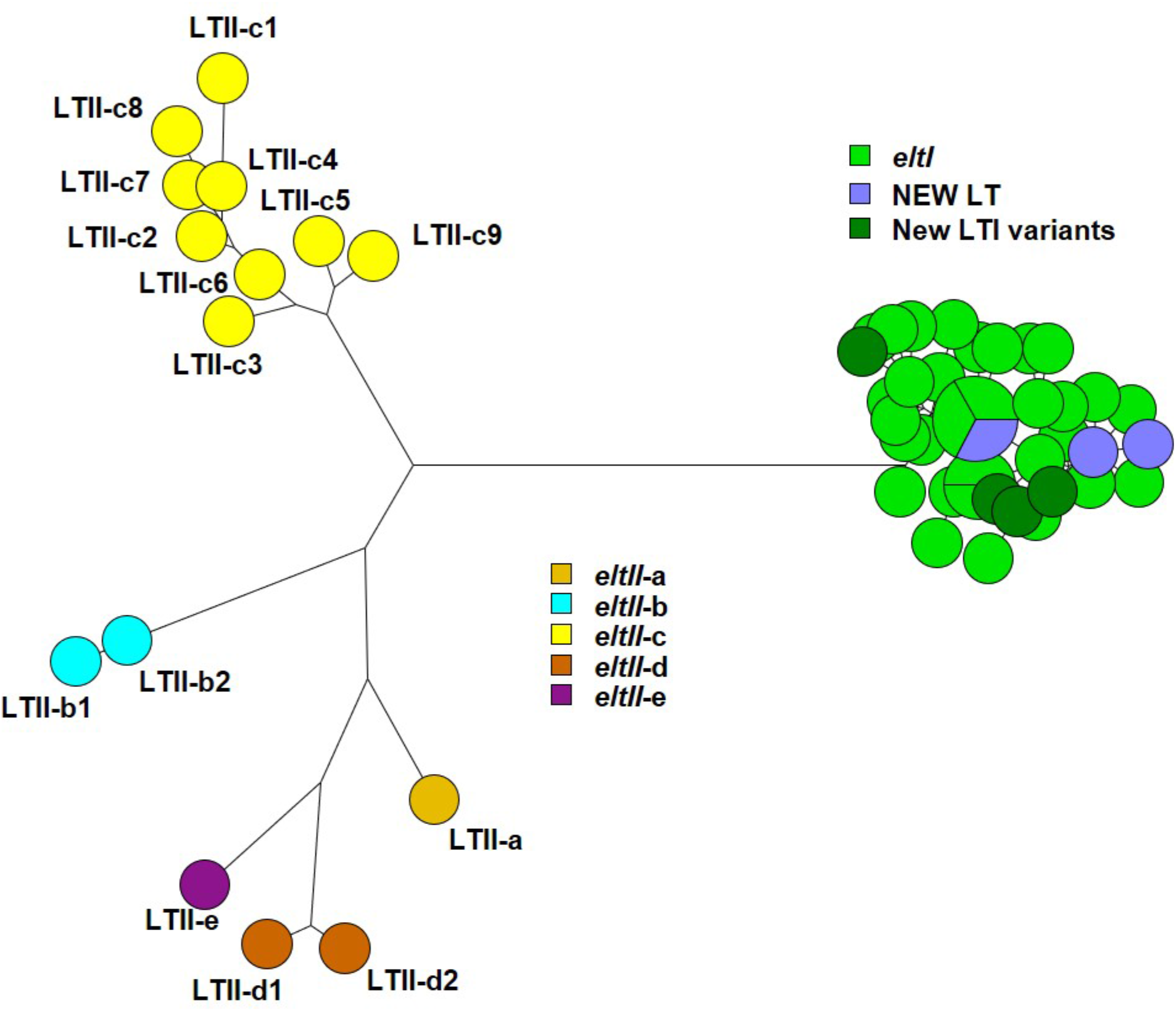
Maximum parsimony tree of 31 LTI and 15 LTII reference nucleotide sequences, three new LTI types (LTI-31, LTI-32 and LTI-33) and four new LTI variants (LTIh-12b, LTIh-15b, LTIh-18b and LTIh-18c) identified in 890 ETEC genomes analyzed with the revised VirulenceFinder database. LTII-a – LTII-e sequences were 48.6 – 54.3% identical to LTI sequences. For a high resolution of *eltI*, see **Figure 3**.

Not all genes were found with 100 % identity, indicating alterations in the sequence compared to the reference sequence found in the database. This could be a result of point mutations or poor sequencing.

### Human-specific ETEC colonization factors

Five-hundred-and-fifty-two (51%) of the genomes used in this study contained genes from two to five different CF types (**Supplementary Table 5)**. Three-hundred-and-thirty-two genomes were positive for only one CF gene (**Supplementary Table 6**). CFAI encoded by *cfaA, cfaB, cfaC, cfaD* and *cfaE* was found in combination with other CFs in 327 genomes (**Supplementary Table 5**). In three ETEC-UPEC_HM_ genomes of serotypes O128ac:H45 (2 genomes) and O153:H46 (1) CFA/I was the only CF (**Supplementary Table 6**). The full gene cluster (*cfaA*, *cfaB*, *cfaC*, *cfaD* and *cfaE* genes) encoding CFA/I was found in two genomes, but in addition to these, 79 genomes had two different *cfaD* genes, all of which were found on two different contigs at identities ranging from 93 % to 100 % and also positive for *cfaE. cfaD* was found once as the only CFA/I gene in 215/330 genomes. *cfaB* and *cfaD* were found in 33/330 genomes. The individual genes *cfaA, cfaB, cfaC, cfaD* and *cfaE* were found in 82, 121, 82, 331 and 82 genomes, respectively. CS2 genes (*cotABCD*) were found in all 42 *cooA* and *cooC* (CS1) sequences but only in four *cooB* and none of the *cooD* sequences. CS2 genes were complete (*cotABCD*) in 36 sequences. CS3 genes were found in 97 sequences, *cstA* (38), *cstB* (62), *cstC* (1), *cstD* (0), *cstE* (0), *cstF* (32), and *cstG* (96 once and one twice). CS4 genes were found in 18 sequences, *csaABCE* (13) and *csaD* (17 once and one twice). CS5 genes were found in 79 sequences, *csfA* (16), *csfB* (78), *csfC* (77), *csfD* (62 once and one twice), *csfE* (3), and *csfF* (55). CS6 genes were found in 240 sequences, *cssA* (237), *cssB* (238), *cssC* (86) and *cssD* (153). The *csvA* gene for CS7 was found in 79 sequences. CS8 genes were found in 115 sequences, *cofBCDEHIJPST* (19), *cofA* (8), *cofF* (9), *cofG* (10), and *cofR* (21). CS12 genes were found in 142 sequences, *cswABCFG* (27), *cswD* (9), *cswE* (18) and *cswR* (142). CS13 genes were found in 11 sequences, *cshADEFG* (11), *cshB* (2) and *cshC* (9). CS14 genes were found in 44 sequences, *csuA* (16 once, 25 twice, one thrice and one four times), *csuA1* (27 once and twice in two), *csuA2* (26 once and twice in one), *csuBD* (44) and *csuC* (42). CS17 genes were found in 27 sequences, *csbAB* (27), *csbC* (26) and *csbD* (one). CS18 genes were found in 15 sequences, *fotABCFGS* (2), *fotD* (1) and *fotT* (15). CS19 genes were found in 17 sequences, *csdAB* (17), *csdC* (14) and *csdD* (3). CS20 genes were found in 27 sequences, *csnA* (11), *csnB* (18), *csnC* (9), *csnD* (13), *csnE* (14) and *csnFG* (27). CS21 genes were found in 255 sequences, *lngBEFGHJPRSTX2* (254; *lngX2* twice in one sequence), *lngC* (26), *lngD* (228), *lngI* (246) and *lngX1* (253). The *cseA* gene for CS22 was found in two sequences. CS23 genes were found in 25 sequences, *aalA* (13), *aalB* (0), *aalCF* (12), *aalD* (2), *aalEG* (10), *aalH* (18) and *aalR* (seven once, two twice and five thrice). CS26 genes were found in 19 sequences, *crsBHST* (5), *crsCFG* (19), *crsD* (1) and *crsE* (18). CS30 genes were found in 32 sequences, *csmAFGST* (32), *csmB* (31), *csmC* (1), *csmD* (12) and *csmE* (20). F17 genes were found in 147 sequences, *F17A* (8), *F17C* (10), *F17D* (141) and *F17G* (14). PCFO71 genes were found in three sequences, *cosABC* (3) and *cosD* (0). Add here that using reads gives another result!

### Animal-specific ETEC colonization factors

F4 genes were found in 129 sequences, *faeA* (74), *faeB* (12), *faeC* (108), *faeD* (30), *faeE* (84), *faeF* (117 once and two sequences twice), *faeGab* (6), *faeGab1* (6), *faeGab2* (0), *faeGac* (79 once and five times twice), *faeGad* (2), *faeG* (0), *faeH* (107 once and one twice), *faeH1* (0), *faeH2* (0), *faeI* (111), *faeI1* (1), *faeI2* (0) and *faeJ* (95 once and one twice). Seven of these 129 were in combination with F5 and *fimF41* (6) and *fimF41* (1), two in combination with F6, one in combination with F18ac; six were F4ab alone, 84 were F4ac alone or ten in combination with F6. Two were F4ad (**Supplementary Table 7**). Sixteen of the 129 F4 genes were found in non-ETEC sequences (**Supplementary Table 7**). F5 was only found in 6 sequences in combination with F4 and *fimF41* genes, *fanABCH* (6), *fanD* (5), *fanE* (0), *fanF* (4) and *fanG* (0). F6 was found in 18 sequences, *fasAB* (17), *fasC* (0), *fasDFG* (16), *fasE* (0) and *fasH* (15). Thirteen of these were in combination with other F genes, F4 (2), F4ac (10) and F18ac (1). F18 genes were found in 33 sequences, *fedAab* (7), *fedAac* (34), *fedAnt* (0), *fedBEF* (33) and *fedC.* Fifteen of these were found in ETEC-ExPEC_JJ_/UPEC_HM_-negative sequences, one in combination with F6 and one ExPEC_JJ_/UPEC_HM_ in combination with F4.

### Combinations of human and animal colonization factor genes

Several putative gene clusters were identified with genes from different colonization factor gene clusters; for example, F4 genes were found in combination with other colonization factor genes. F4-CS23 genes were found in 6 genomes. Additional combinations of F4 and CS23 genes with human and other animal colonization factor genes were found in 16 genomes: specifically, F4-F5-F41-CS23 (6), F4-CS12-CS23 (3), F4-CS12-CS23-CS26 (3), F4-CS12-CS20-CS23 (1), F4-CS12-CS23-CS30 (2) and F4-F17-CS23 (1). Other combinations included F4-F17 (84), F4-F6-F17 (12), F4-CS12-CS13 (6), F4-CS12-CS13-CS26 (2) and F4-CS6-CS12-CS13 (1). The genes were most often located on the same contig indicating that this gene cluster may encode a putative colonization factor. Out of 20 F4-CS23 combinations, fifteen were predicted as ETEC, one as ExPEC_JJ_, one as UPEC_HM_, one as ExPEC_JJ_/UPEC_HM_, and two could not be assigned to a pathotype.colonization

### Additional ETEC non-canonical virulence genes

The genes for enterotoxigenic *E. coli* (ETEC) autotransporter A (*eatA*, three alleles), enterotoxin EspC (*espC*, three alleles) and the plasmid-encoded type II secretion pathway-related protein *etpD* (three alleles) were already included in the original VirulenceFinder database (24). The gene *eatA* was found once in 392 and twice in two genomes, and *espC* was found once in an *eae* positive sequence of *in silico* serotype O71:H49. *etpA, etpB* and *etpC* were found respectively in 31, 340 and 341 sequences. Of note, none of these were also positive for *etpD* found in nine sequences. The adhesin gene *tia* was found once in 182 genomes and twice in one genome. Additionally, genes such as *tibA* and *tibB* were found in 48 and 84 sequences, *tleA* was found twice, and *yghJ* was found once in 899 and twice in 48 sequences.

### Validation of ETECFinder database on the ETEC reference strain H10407

Using the revised database on the annotated sequences for ETEC reference strain H10407 (chromosome, Acc. No. FN649414, and plasmids FN649415.1_p52, FN649415.1_p58, FN649415.1_p666 and FN649415.1_p948) identified *astA, csgA, fdeC, fimH, fyuA, gad, hha, hlyE, irp2, nlpI, shiA, terC, terC, tia, tibA, yehA, yehB, yehC, yehD* and *yghJ* on the chromosome, *anr, astA, cfaA, cfaB, cfaC, cfaD* (two copies)*, cfaE, eatA, estah*-STa2*, etpA, etpB, etpC* on plasmid p948, *eltIAB*-1*, estap*-STa1*, traJ* on plasmid p666 and did not identify any virulence genes on plasmids p52 and p58. The two copies of *cfaD* were both found on plasmid p948 at 100 % identity to Acc. No.s FN649418 (length 375 bp) and M55661 (length 435 bp) at positions 89363..89737, and 43366..43800, respectively. Thus, the genes previously described, *eatA*, heat-stable enterotoxin STa2 (*sta2*), CFA/I fimbriae genes (*cfaABCD*), and the Etp two-partner secretion system and associated glycosyltransferase (*etpBAC*) (25), were all confirmed as being present on plasmid 948.

## DISCUSSION

The decreasing costs of whole genome sequencing (WGS) have led to an increase in bacterial pathogen sequencing. However, extracting the correct data for analysis from WGS data can be challenging, despite its increased availability to routine diagnostic laboratories. To address this issue, we developed, implemented, and evaluated an ETECFinder database to predict ETEC-associated virulence genes based on WGS data. This database has been integrated into the pre-existing VirulenceFinder, which is accessible at www.cge.cbs.dtu.dk/services/VirulenceFinder/. Users can upload preassembled bacterial genomes or short sequence reads, and the CGE webtools have been designed to facilitate use and output for users with limited bioinformatics experience.

Using the revised VirulenceFinder database to analyze 1,083 preassembled genomes, we identified 890 ETEC sequences, while 193 sequences were negative for enterotoxin genes. All sequences had originally been identified as ETEC-positive by PCR, and the failure to detect ETEC genes in 193 genomes could be explained by loss of toxin genes (which are often plasmid-encoded and therefore can be lost during storage) or by poor sequencing and assembly quality. Matches with an identity of 95% or higher are likely due to point mutations or poor sequencing, while an identity score of between 95% and 92% could represent new alleles.

Detecting colonization factors in ETEC is challenging due to their large number, heterogeneity, and lack of standardized tests. However, the detection of LT and ST defines an ETEC isolate, although many such isolates may express colonization factors specific to either animals or humans. In this study, we found 160 sequences with different combinations of human and animal colonization factor genes primarily involving F4 and F18. Further investigation of eight selected genomes revealed that the minor fimbrial subunit encoded by *aalH* (CS23) was located on the same contigs as F4 genes *faeH* and *faeI*. As *aalH* has been described as similar to *faeJ* (97% identity) (26) it could be speculated that some ETEC strains have acquired supplementary genes for the full expression of colonization factors by horizontal gene exchange. While examining such gene exchange in the tested 1,083 sequences is beyond the scope of this study, expanding the VirulenceFinder tool with these ETEC-associated gene alleles will hopefully encourage users to conduct closer analyzes.

Apart from 30 non-ETEC genomes that could be categorized as either ExPEC_JJ_, UPEC_HM_ or both, this study identified four ETEC-ExPEC_JJ_ and one ETEC-UPEC_HM_ genomes. Several studies have identified hybrid STEC-ETEC strains from both humans and animals (1–7). EPEC expressing LT of ETEC has also been described (27). Therefore, it is important to include these ETEC-related genes in a more comprehensive *VirulenceFinder* tool at the CGE website in order to obtain a more complete coverage of the virulence gene repertoire of pathogenic types of *E. coli.*

The standard nomenclature of *est* and *elt* genes is critical to minimize mistakes when analyzing these genes. Current nomenclature is insufficient to classify whether an isolate contains porcine or human-associated *est* genes. Additionally, it is impossible to distinguish between LT types based on the *elt* genes, as the gene name does not refer to the individual LT type. However, the new nomenclature meets these criteria, thus easing analyzes.

We observed several differences in the identification of genes when results obtained by assembled sequences with those obtained by short reads FASTQ files. We would therefore recommend users examine both kinds of sequence data with the *VirulenceFinder* webtool.

## FUTURE PERSPECTIVES

ETEC vaccines are of great importance due to the severity of the infections, primarily in children. A tool such as this could assist in the surveillance of ETEC in order to determine the prevalence of relevant types in different parts of the world, allowing vaccine developers to target the most prevalent types and thus, a more effective vaccine.

