## Supplemental Table 1 for "Construction of the ETECFinder database for the characterisation of enterotoxigenic *Escherichia coli* (ETEC) and revision of the *VirulenceFinder* webtool at the CGE website"

**Supplementary Table 1. Alterations needed after revision of the *VirulenceFinder* database covering both colonization factors and enterotoxins.**

| Gene | Protein function used in the original *VirulenceFinder* database | No. of alleles in the original database | New gene name | New name of the protein function in the revised database | No. of alleles in the revised database |
|---|---|---|---|---|---|
| *cfa_c* | Colonisation factor antigen I | 4 [1] | *cfaA*<br>*cfaB*<br>*cfaC*<br>*cfaE*<br>*cfaD* | CFA/1 Major pilin<br>CFA/1 Major pilin chaperone<br>CFA/1 Usher<br>CFA/1 Pilus length modulator<br>CFA/1 Minor pilin | 2<br>4<br>2<br>1<br>4 |
| *cofA* | Longus type IV pilus subunit | 1 | *cofR*<br>*cofS*<br>*cofT*<br>*cofA*<br>*cofB*<br>*cofC*<br>*cofD*<br>*cofE*<br>*cofF*<br>*cofG*<br>*cofH*<br>*cofI*<br>*cofJ*<br>*cofP* | CS8 Regulator<br>CS8 AraC-like protein<br>CS8 Unknown<br>CS8 Type IV pilin precursor; major pilin<br>CS8 Minor pilin<br>CS8 Unknown<br>CS8 Lipoprotein; secretin<br>CS8 Inner membrane accessory protein (IMAP)<br>CS8 Inner membrane accessory protein (IMAP)<br>CS8 Inner membrane accessory protein (IMAP)<br>CS8 Nucleotide-binding protein; assembly ATPase<br>CS8 Integral membrane protein; inner membrane core protein<br>(IMCP)<br>CS8 Secreted colonization factor<br>CS8 Prepilin peptidase | 2<br>2<br>2<br>2<br>2<br>2<br>2<br>2<br>2<br>2<br>2<br>2<br>2<br>2 |
| *f17-A*<br><br><br>*f17-G* | Subunit A of F17 fimbrial protein<br><br>Adhesin subunit of F17 fimbriae | 7<br><br><br>9 | *F17A*<br>*F17C*<br>*F17D*<br>*F17G* | Major fimbrial subunit of F17<br>F17 Usher<br>F17 Chaperone<br>Minor adhesive subunit of F17 fimbriae | 17<br>11<br>10<br>15 |
| *fanA* | Involved in biogenesis of F5 (K99) fimbriae | 1 | *fanA*<br>*fanB*<br>*fanC*<br>*fanD*<br>*fanE*<br>*fanF* | F5 (K99) Transcriptional regulator<br>F5 (K99) Regulator<br>F5 (K99) Major fimbrial subunit<br>F5 (K99) Usher<br>F5 (K99) Chaperone<br>F5 (K99) Minor fimbrial subunits | 1<br>1<br>3<br>1<br>2<br>1 |

| | | | | | |
|---|---|---|---|---|---|
| | | | *fanG* | F5 (K99) Minor subunit | 1 |
| | | | *fanH* | F5 (K99) Minor subunit | 2 |
| *fasA* | Fimbriae 987P/F6 subunit | 1 | *fasA* | F6 (987P) Major subunit | 1 |
| | | | *fasB* | F6 (987P) Major subunit chaperone | 1 |
| | | | *fasC* | F6 (987P) Adhesin-specific chaperone | 1 |
| | | | *fasD* | F6 (987P) Outer membrane usher | 1 |
| | | | *fasE* | F6 (987P) Chaperone-like | 1 |
| | | | *fasF* | F6 (987P) Minor subunit linker | 1 |
| | | | *fasG* | F6 (987P) Minor adhesive subunit | 1 |
| | | | *fasH* | F6 (987P) Positive regulator, previously *fapR* (8) | 1 |
| *fedA* | Fimbrial F107 subunit A | 3 | *fedAab* | F18 (F107) Major serotype specific fimbrial subunit serotype ab | 5 |
| *fedF* | Fimbrial adhesin AC precursor | | *fedAac* | F18 (F107) Major serotype specific fimbrial subunit serotype ac | 11 |
| | | | *fedAnt* | F18 (F107) Major serotype specific fimbrial subunit serotype nt | 1 |
| | | | *fedB* | F18 (F107) Usher | 2 |
| | | | *fedC* | F18 (F107) Chaperone | 1 |
| | | | *fedE* | F18 (F107) Minor fimbrial subunits | 3 |
| | | 6 | *fedF* | F18 (F107) Minor adhesive subunit | 12 |
| *K88ab* | K88/F4 subunit [2] | 10 | *faeA* | F4 Regulator | 2 |
| | | | *faeB* | F4 Partial | 2 |
| | | | *faeC* | F4 Minor fimbrial subunits | 4 |
| | | | *faeD* | F4 Usher | 8 |
| | | | *faeE* | F4 Chaperone | 9 |
| | | | *faeF* | F4 Minor fimbrial subunit | 8 |
| | | | *faeGab* | F4ab, Major serotype specific fimbrial subunit | 3 |
| | | | *faeGac* | F4ac | 14 |
| | | | *faeGad* | F4ad | 1 |
| | | | *faeGW* | F4W | 1 |
| | | | *faeH* | F4 Fimbrial minor subunit | 8 |
| | | | *faeI* | F4 Fimbrial minor subunit | 17 |
| | | | *faeJ* | F4 Fimbrial minor subunit | 5 |
| *fimF41* | Mature Fim41a/F41 protein | 2 | *fim41* | FimF41 Major fimbrial subunit | 4 |
| *lngA* | Longus type IV pilus | | *lngX1* | - | 1 |
| | | | *lngR* | CS21 Putative *papB*-like regulator | 2 |
| | | | *lngS* | CS21 Putative AraC-like regulator | 3 |
| | | | *lngT* | - | 1 |
| | | | *lngX2* | - | 3 |
| | | 2 | *lngA* | CS21 Major pilin subunit | 3 |
| | | | *lngB* | CS21 Minor pilin subunit | 1 |

| | | | | | |
|---|---|---|---|---|---|
| | | | *lngC* | - | 1 |
| | | | *lngD* | CS21 Putative outer membrane protein | 3 |
| | | | *lngE* | CS21 Putative inner membrane protein | 1 |
| | | | *lngF* | CS21 Putative inner membrane protein | 1 |
| | | | *lngG* | CS21 Putative periplasmic protein | 2 |
| | | | *lngH* | CS21 Putative nucleotide-binding protein | 2 |
| | | | *lngI* | CS21 Putative inner membrane protein | 2 |
| | | | *lngJ* | CS21 Putative ATPase-like | 3 |
| | | | *lngP* | CS21 Putative prepilin peptidase | 1 |
| *ltcA* [3] | Heat-labile enterotoxin A subunit | 17 | *eltI*AB_*1-30* | Heat-labile enterotoxin holotoxin AB subunit sequences; LTIh1-LTIh30 [4] | 31 |
| | | | *eltI*ABp | LTIp | 1 |
| *sta1* | Heat-stabile enterotoxin ST-Ia | 2 | *estap* | Heat-stable enterotoxin STa1, STa4, STa5, STa6, STa7, STa8 and STa9 porcine variants | 15 [5] |
| | | | *estah* | Heat-stable enterotoxin STa2 and STa3 human variants | 4 |
| *stb* | Heat-stabile enterotoxin II | 3 | *estb* | Heat-stable enterotoxin STb1 porcine variants | 3 |
| | | | | Heat-stable enterotoxin STb2 human variant | 1 |

[1] Two *cfaC* alleles in the original database were actually *csaA* (CS4) and *csuA* (CS14) respectively

[2] Seven of these were actually F4ac, one F4ad and one F4W.

[3] The original designation *ltcA* (“c” for chicken) was deleted. The revision included full holotoxin *eltIAB* sequences for 31 LTI types.

[4] Three different nucleotide alleles had synonymous nucleotide substitutions: Acc. JX504011 = CP000795, EU113242 = EU113244, and EU113244 = EU113245.

[5] Two different STa4 alleles that had synonymous nucleotide substitutions: Acc. CP012501 = KT992796.
