## Supplemental Table 2 for "Construction of the ETECFinder database for the characterisation of enterotoxigenic *Escherichia coli* (ETEC) and revision of the *VirulenceFinder* webtool at the CGE website"

**Supplementary Table 2.** LT types based on the specific combination of the A and B subunit.

| LT-Type | A subunit | B subunit(s) |
|---|---|---|
| LT1 | 1 | 1, 10, 12, 13, 18, 21, 23, 24, 25, 26, 27, 28 |
| LT1p | 1p, 4, 6 | |
| LT2 | 2 | 2, 7, 15, 16, 22 |
| LT3 | 3, 5 | 3, 8 |
| LT4 | 1p, 4, 6 | 4 |
| LT5 | 3, 5 | 5 |
| LT6 | 1p, 4, 6 | 6 |
| LT7 | 7 | 2, 7, 15, 16, 22 |
| LT8 | 8 | 3, 8 |
| LT9 | 9 | 9, 11, 19 |
| LT10 | 10 | 1, 10, 12, 13, 18, 21, 23, 24, 25, 26, 27, 28 |
| LT11 | 11 | 9, 11, 19 |
| LT12 | 12 | 1, 10, 12, 13, 18, 21, 23, 24, 25, 26, 27, 28 |
| LT13 | 13 | 1, 10, 12, 13, 18, 21, 23, 24, 25, 26, 27, 28 |
| LT14 | 14 | 14 |
| LT15 | 15 | 2, 7, 15, 16, 22 |
| LT16 | 16 | 2, 7, 15, 16, 22 |
| LT17 | 17 | 17, 20, 29, 30 |
| LT18 | 18 | 1, 10, 12, 13, 18, 21, 23, 24, 25, 26, 27, 28 |
| LT19 | 19 | 9, 11, 19 |
| LT20 | 20 | 17, 20, 29, 30 |
| LT21 | 21 | 1, 10, 12, 13, 18, 21, 23, 24, 25, 26, 27, 28 |
| LT22 | 22 | 2, 7, 15, 16, 22 |
| LT23 | 23 | 1, 10, 12, 13, 18, 21, 23, 24, 25, 26, 27, 28 |
| LT24 | 24 | 1, 10, 12, 13, 18, 21, 23, 24, 25, 26, 27, 28 |
| LT25 | 25 | 1, 10, 12, 13, 18, 21, 23, 24, 25, 26, 27, 28 |
| LT26 | 26 | 1, 10, 12, 13, 18, 21, 23, 24, 25, 26, 27, 28 |
| LT27 | 27 | 1, 10, 12, 13, 18, 21, 23, 24, 25, 26, 27, 28 |
| LT28 | 28 | 1, 10, 12, 13, 18, 21, 23, 24, 25, 26, 27, 28 |
| LT29 | 29, 30 | 17, 20, 29, 30 |
| LT30 | 29, 30 | 17, 20, 29, 30 |
