## Supplemental Table 3 for "Construction of the ETECFinder database for the characterisation of enterotoxigenic *Escherichia coli* (ETEC) and revision of the *VirulenceFinder* webtool at the CGE website"

**Supplementary Table 3.** ETEC associated genes including CF type or protein name, and number of alleles added as new to the *VirulenceFinder* database.

| Gene | Description | CF type or name of protein | No. of alleles |
|---|---|---|---|
| *cooA* | Major subunit | CS1 | 1 |
| *cooB* | Chaperone | CS1 | 1 |
| *cooD* | Chaperone | CS1 | 1 |
| *cooC* | Usher | CS1 | 1 |
| *cotB* | Major pilin chaperone | CS2 | 1 |
| *cotA* | Major pilin chaperone | CS2 | 1 |
| *cotC* | Usher | CS2 | 1 |
| *cotD* | Chaperone | CS2 | 1 |
| *cstA* | Major pilin chaperone | CS3 | 1 |
| *cstBCDEF* | Outer membrane usher | CS3 | 6 [1] |
| *cstG* | Major pilin | CS3 | 3 |
| *cstH* | Major pilin reported (1) but not found | CS3 | - |
| *csaA* | Periplasmic chaperone-like protein | CS4 | 1 |
| *csaB* | CS4 major fimbriae subunit | CS4 | 1 |
| *csaC* | Usher protein | CS4 | 1 |
| *csaE* | Tip-associated protein | CS4 | 1 |
| *csaD*’ | Deleted regulatory protein | CS4 | 1 |
| *csfA* | major subunit | CS5 | 1 |
| *csfB* | coli surface factor five B, Chaperone | CS5 | 1 |
| *csfC* | Outer membrane usher protein | CS5 | 1 |
| *csfE* | Pilus length modulator | CS5 | 1 |
| *csfF* | Minor pilin chaperone | CS5 | 1 |
| *csfD* | Minor fimbrial subunit | CS5 | 1 |
| *cssA* | Structural subunit A | CS6 | 4 |
| *cssB* | Structural subunit B | CS6 | 2 |
| *cssC* | Periplasmic chaperone | CS6 | 8 |
| *cssD* | Usher protein | CS6 | 4 |
| *csvA* | Major subunit | CS7 | 1 |
| *cswA* | Major subunit | CS12 | 1 |
| *cswB* [2] | Chaperone | CS12 | 1 |
| *cswC* | Chaperone | CS12 | 1 |
| *cswD* | Outer membrane usher | CS12 | 1 |
| *cswE* | Chaperone | CS12 | 1 |
| *cswF* | Minor subunit | CS12 | 1 |
| *cswG* | Putative adhesin | CS12 | 1 |
| *cswR* | Transcriptional activator | CS12 | 1 |
| *cshA* | Minor pilin | CS13 | 1 |

| | | | |
|---|---|---|---|
| *cshB* | Usher | CS13 | 1 |
| *cshC* | Chaperone | CS13 | 1 |
| *cshD* | Minor pilin | CS13 | 1 |
| *cshE* | Pilin | CS13 | 1 |
| *cshF* | Minor pilin | CS13 | 1 |
| *cshG* | Minor pilin | CS13 | 1 |
| *csuA* | Major fimbrial subunit | CS14 | 3 |
| *csuB* | Periplasmic chaperone | CS14 | 1 |
| *csuC* | Outer membrane usher | CS14 | 1 |
| *csuD* | Minor fimbrial subunit | CS14 | 1 |
| *nfaA* | Nonfimbrial adhesin A | CS15 | 1 |
| *csbB* | Chaperone | CS17 | 2 |
| *csbA* | Fimbriae major subunit | CS17 | 1 |
| *csbC* | Usher | CS17 | 1 |
| *csbD* | Assembly protein/minor subunit | CS17 | 2 |
| *fotA* | Major subunit | CS18 | 2 |
| *fotB* | Major chaperone | CS18 | 1 |
| *fotC* | Minor chaperone | CS18 | 1 |
| *fotD* | Usher | CS18 | 1 |
| *fotE* | Minor chaperone | CS18 | 1 |
| *fotF* | Minor subunit | CS18 | 1 |
| *fotG* | Minor subunit-adhesin | CS18 | 1 |
| *fotS* | Site-specific recombinase | CS18 | 1 |
| *fotT* | Site-specific recombinase | CS18 | 1 |
| *csdB* | Periplasmic chaperone | CS19 | 1 |
| *csdA* | Major fimbrial subunit | CS19 | 1 |
| *csdC* | Outer membrane usher | CS19 | 1 |
| *csdD* | Minor fimbrial subunit | CS19 | 1 |
| *csnA* | Fimbria major subunit | CS20 | 2 |
| *csnB* [2] | Putative major subunit chaperone | CS20 | 2 |
| *csnC* | Putative adhesin-specific chaperone | CS20 | 2 |
| *csnD* | Putative outer membrane fimbrial usher | CS20 | 2 |
| *csnE* | Putative chaperone-like | CS20 | 1 |
| *csnF* | Putative minor subunit | CS20 | 2 |
| *csnG* | Putative minor adhesive tip subunit | CS20 | 2 |
| *cseA* | Adhesin protein | CS22 | 1 |
| *aalR* | Putative transcriptional regulator | CS23 | 1 |
| *aalA* | Minor structural subunit | CS23 | 1 |
| *aalB* | Usher | CS23 | 1 |
| *aalC* | Chaperone | CS23 | 1 |

| | | | |
|---|---|---|---|
| *aalD* | Minor structural subunit | CS23 | 1 |
| *aalE* | Major structural subunit | CS23 | 1 |
| *aalF* | Minor structural subunit | CS23 | 1 |
| *aalG* | Minor structural subunit | CS23 | 1 |
| *aalH* | Minor structural subunit | CS23 | 1 |
| *crsB* | Molecular chaperone | CS26 | 1 |
| *crsC* | Fimbria/pilus periplasmic chaperone | CS26 | 1 |
| *crsD* | Fimbrial biogenesis outer membrane usher protein | CS26 | 1 |
| *crsE* | Fimbria/pilus periplasmic chaperone | CS26 | 1 |
| *crsF* | Fimbrial protein | CS26 | 1 |
| *crsG* | Hypothetical protein | CS26 | 1 |
| *crsH* | Coli surface antigen | CS26 | 2 |
| *crsS* | Tyrosine-type DNA invertase | CS26 | 1 |
| *crsT* | Tyrosine-type DNA invertase | CS26 | 1 |
| *cmaHa* | Coli surface antigen a | CS27A | 1 |
| *cmaHb* | Coli surface antigen b | CS27B | 1 |
| *cnmHa* | Coli surface antigen a | CS28A | 1 |
| *cnmHb* | Coli surface antigen b | CS28B | 1 |
| *csmS* | Site specific recombinase | CS30 | 1 |
| *csmT* | Site specific recombinase | CS30 | 1 |
| *csmA* | Major subunit | CS30 | 1 |
| *csmB* | Chaperone | CS30 | 1 |
| *csmC* | Chaperone | CS30 | 1 |
| *csmD* | Usher | CS30 | 1 |
| *csmE* | Chaperone | CS30 | 1 |
| *csmF* | Minor subunit | CS30 | 1 |
| *csmG* | Fimbrial adhesin | CS30 | 1 |
| *cosA* | Major fimbrial subunit | PCFO71 | 1 |
| *cosB* | Periplasmic chaperone | PCFO71 | 1 |
| *cosC* | Outer membrane usher | PCFO71 | 1 |
| *cosD* | Minor fimbrial subunit | PCFO71 | 1 |
| *Tia* | Invasion determinant | Tia | 9 |
| *tibA* | Adhesin/invasin (Glycoprotein) | TibA | 4 |
| *tibC* | Glycosyltransferase | TibC | 7 |
| *etpA* | Invasin and adherence | EtpA | 19 |
| *etpB* | Nonfimbrial adhesin/TPS transporter | EtpB | 4 |
| *etpC* [4] | Glycotransferase | EtpC | 9 |
| *tleA* | *tsh*-like ETEC autotransporters/Glycosyltransferase | TleA | 2 |
| *yghJ* | Metalloprotease | YghJ | 5 |

[1] *cstB* contains 11 ORFs of which five are considered to be functional usher genes *cstB* (2 alleles), *cstC*, *cstD*, *cstE*, and *cstF* (one each)

[2] *cswB* (CS12) is identical to *csnB* (CS20)
