## Supplemental Table 4 for "Construction of the ETECFinder database for the characterisation of enterotoxigenic *Escherichia coli* (ETEC) and revision of the *VirulenceFinder* webtool at the CGE website"

**Supplementary Table 4.** Description of enterotoxins and the alleles as well as their gene names. Changes to the current database are indicated.

| LT-type | ST-type | SUM |
|---|---|---|
| **LT1b** | *estap*; STa1-01 | 14 |
| **LT3** | - | 6 |
| **LT7** | - | 2 |
| **LT8** | - | 14 |
| **LT11** | - | 10 |
| **LT12** | - | 2 |
| **LT12+LT13+LT25** | - | 1 |
| **LT12b** | - | 2 |
| **LT13** | - | 17 |
| **LT15** | *estah*; STa3-01 | 39 |
| **LT15** | *estah;* STa2-01 | 19 |
| **LT15** | *estah;* STa3-05 | 3 |
| **LT15** | - | 23 |
| **LT15b** | - | 11 |
| **LT17** | - | 25 |
| **LT18** | *estap*; STa1-02 | 21 |
| **LT18** | *estap*; STa1-02; *estah*; STa3-02 | 8 |
| **LT18** | *estap*; STa1-02b | 1 |
| **LT18** | - | 2 |
| **LT18b** | *estap*; STa1-02 | 6 |
| **LT18c** | *estap*; STa1-01 | 1 |
| **LT19** | *estap*; STa1-01 | 1 |
| **LT20** | *estah*; STa3-01 | 3 |
| **LT21** | - | 1 |
| **LT22** | - | 1 |
| **LT23** | *estap*; STa1-01 | 12 |
| **LT24** | *estap*; STa1-01 | 12 |
| **LT25** | *estap*; STa1-01 | 6 |
| **LT26** | *estap*; STa7-01 | 1 |
| **LT27** | *estap*; STa1-01 | 1 |
| **LT28** | *estap*; STa1-01 | 1 |
| **LT29** | *estah*; STa3-01 | 3 |
| **LT30** | *estah*; STa3-01 | 90 |
| **LT30** | *estap*; STa1-01 | 1 |
| **New LT31** | *estap*; STa1-01 | 1 |
| **New LT32** | *estap*; STa1-01 | 1 |
| **New LT32** | *estap*; STa1-05 | 14 |
| **New LT32** | - | 51 |
| **New LT33** | *estap*; STa1-01 | 15 |
| New LT33 | - | 12 |
| **LTp1 (previous LT4)** | *estap*; STa1-01, *estb*; STb1-01 | 13 |
| **LTp1 (previous LT4)** | *estap*; STa1-01, *estb*; STb1-03 | 4 |

| | | |
|---|---|---|
| **LTp1 (previous LT4)** | *estb*; STb1-01 | 107 |
| **LTp1 (previous LT4)** | *estb*; STb1-01-NEW | 1 |
| **LTp1 (previous LT4)** | - | 3 |
| - | *estah*; STa3-01 | 88 |
| - | *estah*; STa3-02 | 38 |
| - | *estah;* STa2-01 | 63 |
| - | *estah;* STa2-02 | 1 |
| - | *estah;* STa3-04 | 1 |
| - | *estah;* STa3-06 | 2 |
| - | *estap*; STa1, *estb*; STb1-03 | 1 |
| - | *estap*; STa1-01 | 17 |
| - | *estap*; STa1-01, *estb*; STb1-01 | 1 |
| - | *estap*; STa1-01, *estb*; STb1-03 | 11 |
| - | *estap*; STa1-03 | 78 |
| - | *estap*; STa1-06 | 1 |
| - | *estap*; STa4-04 | 2 |
| - | *estap*; STa5-07 | 1 |
| - | *estb*; STb1-01 | 1 |
| - | - | 193 |
| **Total** | | **1083** |
