## Supplemental Table 5 for "Construction of the ETECFinder database for the characterisation of enterotoxigenic *Escherichia coli* (ETEC) and revision of the *VirulenceFinder* webtool at the CGE website"

**Supplementary Table 5.** Combinations of CF genes found in 441 ETEC and 11 non-ETEC sequences. Presence of one gene specific for a CF is enough to be reported, i.e., all genes encoding a CF may not be present in all CF combinations.

| Combinations | ETEC | Non-ETEC | Total |
|---|---|---|---|
| CFA/I; CS1 | | 2 | 2 |
| CFA/I; CS1; CS21 | | 3 | 3 |
| CFA/I; CS1; CS3 | 1 | | 1 |
| CFA/I; CS1; CS3; CS21 | 36 | | 36 |
| CFA/I; CS12 | 1 | | 1 |
| CFA/I; CS12; CS21 | 12 | | 12 |
| CFA/I; CS14 | 43 | | 43 |
| CFA/I; CS2; CS21 | 1 | | 1 |
| CFA/I; CS2; CS3 | 2 | | 2 |
| CFA/I; CS2; CS3; CS21 | 30 | | 30 |
| CFA/I; CS2; CS3; CS21; CS23 | 1 | | 1 |
| CFA/I; CS21 | 77 | | 77 |
| CFA/I; CS3 | 6 | | 6 |
| CFA/I; CS3; CS21 | 19 | | 19 |
| CFA/I; CS4; CS21 | 2 | 1 | 3 |
| CFA/I; CS4; CS6 | 12 | | 12 |
| CFA/I; CS6 | 75 | 2 | 77 |
| CFA/I; CS6; CS23 | 3 | | 3 |
| CFA/I; CS6; CS8 | 6 | | 6 |
| CS12; CS13 | 6 | | 6 |
| CS12; CS13; CS26 | | 2 | 2 |
| CS12; CS13; CS30 | 2 | | 2 |
| CS12; CS20 | 21 | | 21 |
| CS12; CS23 | 3 | | 3 |
| CS12; CS23; CS26 | 3 | | 3 |
| CS12; CS26 | 14 | | 14 |
| CS12; CS2CS23 | 1 | | 1 |
| CS17; CS17 | 27 | | 27 |
| CS18; CS30 | 13 | | 13 |
| CS2; CS21 | | 1 | 1 |
| CS2; CS3; CS21 | 1 | | 1 |
| CS23; F17 | 1 | | 1 |
| CS4; CS6 | 2 | | 2 |
| CS5; CS6; CS7 | 56 | | 56 |
| CS5; CS7 | 23 | | 23 |
| CS6; CS12 | 18 | | 18 |
| CS6; CS12; CS13 | 1 | | 1 |
| CS6; CS12; CS18 | 1 | | 1 |
| CS6; CS21 | 39 | | 39 |
| CS6; CS8 | 8 | | 8 |
| CS8; F17 | 1 | | 1 |
| **Total:** | **441** | **11** | **552** |
