## Supplemental Table 6 for "Construction of the ETECFinder database for the characterisation of enterotoxigenic *Escherichia coli* (ETEC) and revision of the *VirulenceFinder* webtool at the CGE website"

**Supplementary Table 6.** Single CF genes found in 297 ETEC, one $UPEC_{HM}$, one ETEC-$UPEC_{HM}$, four ETEC-$ExPEC_{JJ}$, six $ExPEC_{JJ}$/$UPEC_{HM}$, one $ExPEC_{JJ}$, and 22 ETEC-$ExPEC_{JJ}$/$UPEC_{HM}$-negative sequences.

| CF type single | $ExPEC_{JJ}$ | $ExPEC_{JJ}$ /$UPEC_{HM}$ | negative | $UPEC_{HM}$ | ETEC/ $ExPEC_{JJ}$ | ETEC | ETEC/$UPEC_{HM}$ | SUM |
|---|---|---|---|---|---|---|---|---|
| F17 | | | 8 | | | 136 | 1 | 144 |
| CS12 | | | 1 | | | 56 | | 57 |
| CS21 | | | 3 | | | 29 | | 32 |
| CS17 | | | | | | 27 | | 27 |
| CS30 | | | | | | 17 | | 17 |
| CS6 | | | 3 | | 4 | 10 | | 17 |
| CS23 | 1 | 1 | 3 | 1 | | 7 | | 13 |
| CS8 | | | 1 | | | 6 | | 7 |
| CS20 | | 5 | | | | | | 5 |
| PCFO71 | | | | | | 3 | | 3 |
| CFAI [1)] | | | | | | 3 | | 3 |
| CS22 | | | 2 | | | | | 2 |
| CS14 | | | | | | 1 | | 1 |
| CS4 | | | | | | 1 | | 1 |
| CS18 | | | | | | 1 | | 1 |
| CS3 | | | 1 | | | | | 1 |
| **Total** | **1** | **6** | **22** | **1** | **4** | **297** | **1** | **332** |

[1)] Found in serotypes O128ac:H45 (2 genomes) and O153:H46 (1).
