## Supplemental Table 7 for "Construction of the ETECFinder database for the characterisation of enterotoxigenic *Escherichia coli* (ETEC) and revision of the *VirulenceFinder* webtool at the CGE website"

**Supplementary Table 7.** Animal fimbriae found in 193 sequences.

| F type | number |
|---|---|
| F4 [1), 2), 3), 4)] | 28 |
| F4ab | 6 |
| F4ac | 84 |
| F4ad | 2 |
| F6 | 5 |
| F4; fimF41 [2)] | 1 |
| F4; F5; fimF41 | 6 |
| F4; F6 | 2 |
| F4ac; F6 | 10 |
| F4; F18 [2)] | 1 |
| F6; F18ac | 1 |
| F18 [1)] | 15 |
| F18ab | 7 [5)] |
| F18ac | 25 |
| Total | 193 |

[1)] Eleven and 15 ETEC-$ExPEC_{JJ}$/$UPEC_{HM}$-negative sequences were positive for F4 and F18 respectively.

[2)] Three $ExPEC_{JJ}$/$UPEC_{HM}$ sequences positive for F4, F4/F18 and F4/*fimF41* respectively

[3)] One $ExPEC_{JJ}$ for F4.

[4)] One $UPEC_{HM}$ was positive for F4

[5)] All seven were positive for *stx2e*-O139-S1191 (Acc. No. M21534).
